# Cargo-Loading of Misfolded Proteins into Extracellular Vesicles: The CSPα-EV Export Pathway

**DOI:** 10.1101/310219

**Authors:** Desmond Pink, Julien Donnelier, John Lewis, Janice E.A. Braun

## Abstract

Extracellular vesicles (EVs) are secreted vesicles of diverse size and cargo that are implicated in the cell-to-cell transmission of disease-causing-proteins in several neurodegenerative diseases. Mutant huntingtin, the disease-causing entity in Huntington’s disease, has an expanded polyglutamine track at the N terminus that causes the protein to misfold and form toxic intracellular aggregates. In Huntington’s disease, mutant huntingtin aggregates are transferred between cells by an unknown route. We have previously identified a cellular pathway that is responsible for the export of mutant huntingtin via extracellular vesicles, given the heterogeneity of EVs, here we examine the specific EVs involved. In this work we expressed a form of polyglutamine expanded huntingtin (GFP-tagged 72Qhuntingtin^exon1^) in cells to assess the EVs involved in cellular export. We demonstrate that the molecular chaperone, cysteine string protein (CSPα; DnaJC5), mediates export of disease-causing-polyglutamine-expanded huntingtin cargo via two distinct vesicle populations of 180-240nm and 15-30μm. In doing so, our data links the molecular chaperone, CSPα, and the packaging of pathogenic misfolded huntingtin into two separate extracellular vesicles pathways.

## Introduction

The cell-to-cell transfer of extracellular vesicles (EVs) is a conserved process. *In vivo*, the continuous exchange among different cells generates a dynamic and heterogeneous pool of EVs (Pegtel and Gould, 2019). EVs come in different sizes and carry different cargoes that exert profound effects in recipient cells following uptake. The physiological roles of EVs include exchanging physiological information between cells as well as removing unwanted proteins from cells (Hill, 2019; Pegtel and Gould, 2019; Vella et al., 2016). EVs are also implicated in disease progression however their role is far from clear. How EVs facilitate the spread of disease in cancer and neurodegenerative disease and what distinguishes physiological from pathological EVs is a current focus of investigation (Hill, 2019; Pegtel and Gould, 2019). While complex cargoes of DNA, RNA, proteins, lipids and metabolites are packaged in EVs for delivery to recipient cells, our understanding of the mechanisms that target proteins to EVs is rudimentary in comparison to conventional secretion.

Trinucleotide repeat expansions of the huntingtin gene cause Huntington’s disease, a progressive neurodegenerative disorder that manifests in midlife (Gusella and MacDonald, 2000). Aggregates of polyglutamine-expanded huntingtin are found within genetically normal tissue grafted into patients with progressing Huntington’s Disease, indicating cell-to-cell transit of huntingtin aggregates *in vivo* (Cicchetti et al., 2014). To date, the majority of studies conducted to evaluate mechanisms that suppress aggregation of polyQ expanded proteins like mutant huntingtin have focused on the intracellular activities of molecular chaperones. In this work we have chosen to focus on the cellular export of aggregated huntingtin in a cell culture model. We previously reported that cysteine string protein (CSPα; DnaJC5), an abundant molecular co-chaperone critical for the maintenance of synapses (Fernandez-Chacon et al., 2004; Zinsmaier et al., 1994) facilitates mutant huntingtin export. CSPα is a member of the J protein family, key regulators of the cellular Hsp70 machinery. The human genome encodes 53 J proteins that deliver misfolded protein substrates to Hsp70 and activate Hsp70ATPase activity through a conserved histidine, proline, aspartate (HPD) motif. Here we confirm and extend our previous findings that CSPα exports mutant huntingtin in EVs. Using nanoscale flow cytometry and live cell imaging we identify two subpopulations of EVs, 180-240nm and 15-30μm, responsible for the CSPα-mediated export of mutant huntingtin. Our data highlight the active role EV export plays in the cellular protein quality control network in general and provides strong evidence that CSPα specifically influences Huntington Disease progression.

## Results

### CSPα EV export of 180-240nm EVs

We first asked the question; Is misfolded huntingtin cargo common to all EVs or to select EVs? To address this question, media from CAD neural cells expressing CSPα and GFP-tagged 72Q huntingtin^exon1^ was collected and centrifuged at 300Xg for 5 min to remove cell debris and EVs evaluated by nanoscale flow cytometry. Figure 1 shows that secretion of EVs containing GFP-tagged 72Q huntingtin^exon1^ is dramatically increased when cells express CSPα (p < 0.0001), confirming our previous work. As anticipated, EVs released from CAD cells in the presence of CSPα are heterogeneous (Figure 2A). Small <180nm EVs, presumably exosomes, comprised the majority of exported vesicles (81.6%). However, the bulk of EVs packaged with mutant huntingtin fall within the 180-240nm range (Figure 2A). The 180-240nm EVs make up 17% of the total EVs (Figure 2A and supplementary Figure 1) and CSPα increased export of EVs containing GFP-tagged 72Q huntingtin^exon1^ within the 180-240nm EV subpopulation by >400%.

**Figure 1.**
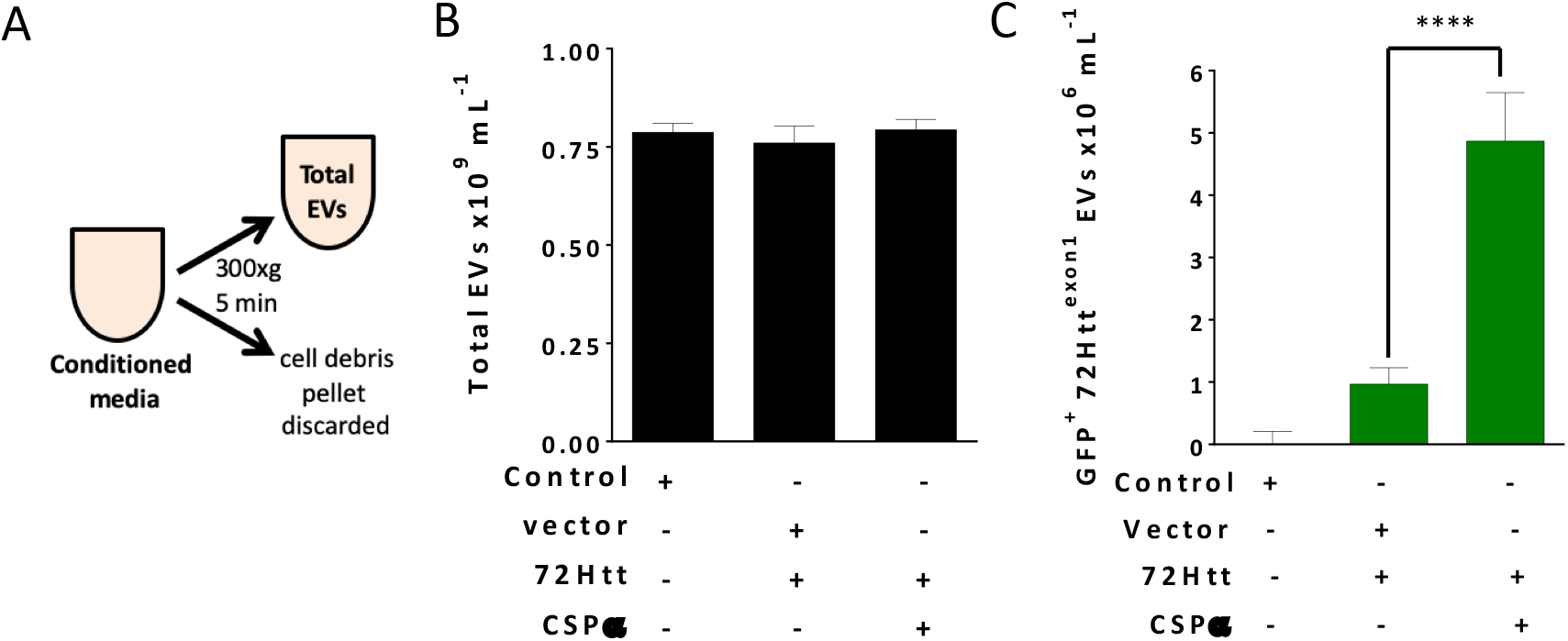
Nanoscale Flow Cytometry Showing the Influence of CSPα on EV Export. (A) Experimental approach, EVs were examined without enrichment. (B) Bar graph of the concentration of total EVs and (C) GFP-72Htt^exon1^-containing EVs as determined by nanoscale flow cytometry. (****P<0.0001)

**Figure 2.**
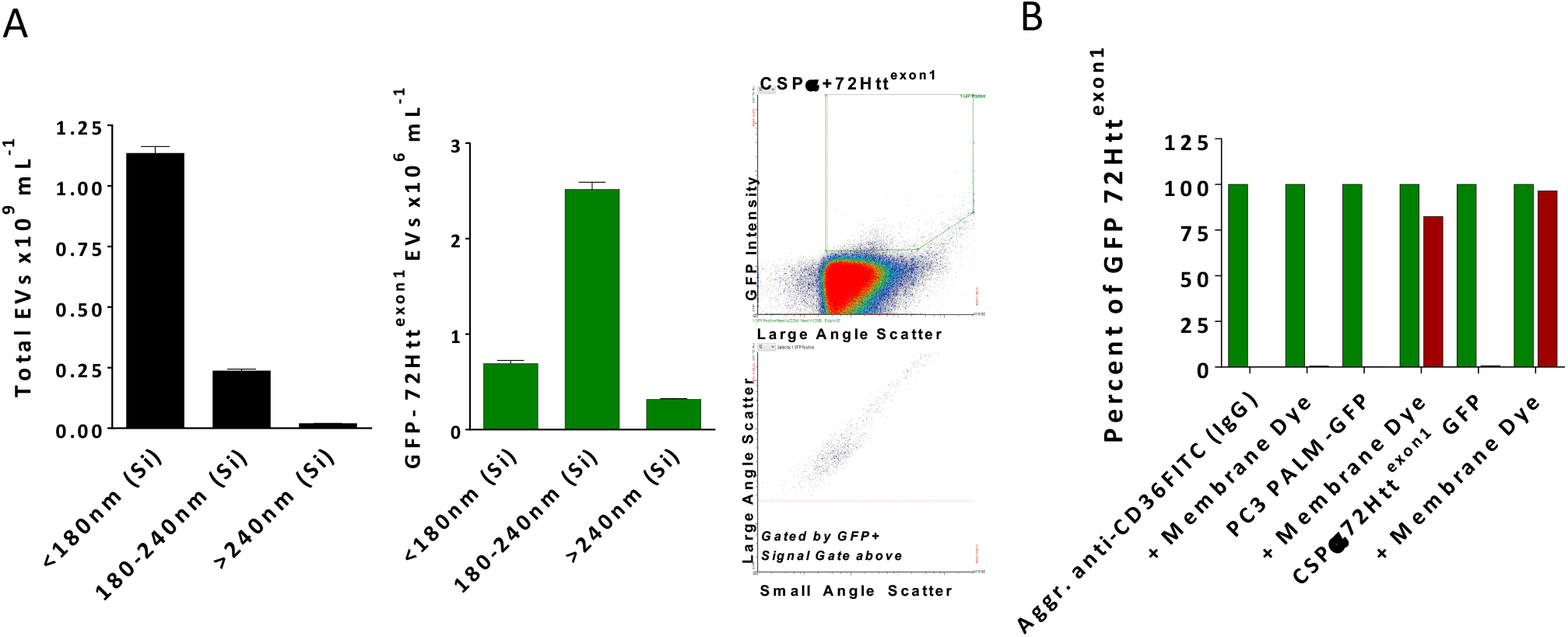
CSPα export of 180-240nm EVs. (A) Bar graph showing the sizes of the total EVs (left panel) and GFP-72Htt^exon1^ containing EVs (middle panel) Silica (Si). The right panel shows the flow cytometry scatter plot. (B) Aggregated anti-CD36 FITC, EVs containing PC3 Palm-GFP and EVs containing GFP-72Htt^exon1^ were stained with Deep Red membrane dye. The experiment was replicated ten-twelve times.

We next sought to determine if any of the GFP particles detected by nanoscale flow cytometry were free floating GFP-tagged 72Q huntingtin^exon1^ entities. EVs were labeled with Cell Mask Deep Red plasma membrane stain for 30 min at 37°C. 95% of GFP-72Q huntingtin^exon1^ particles were found to costain with Deep Red stain, demonstrating the GFP particles are membrane bound vesicles and not free-floating entities (Figure 2B and supplementary Figure 2). By comparison, control protein aggregates of anti-CD36 did not stain with Deep Red (0.6%) and EVs containing the control protein PC3 Palm-GFP were Deep Red positive (80%). The 180-240nm EVs containing exported huntingtin were detergent-sensitive (data not shown), as anticipated. Together, these results show that CSPα expression correlates with an increase in export of GFP-72Q huntingtin^exon1^ in vesicles that are 180-240nm in diameter.

### CSPα EV export of 15-30μm EVs

We also observed mutant huntingtin in large EVs outside of the ~100nm-1300nm range of nanoscale flow cytometry. Figure 3A is a live cell imaging time lapse showing representative export of a large EV containing GFP-72Q huntingtin^exon1^ cargo from CAD cells reminiscent of neural exophers. To further examine these larger EVs, media was collected from CAD cells and submitted to sequential (70μm (nylon) 40μm/1μm (PET Polyethylenterephthalat) filtration (Figure 3B). Despite their size – large EVs were found to be pliant and pass through filters. Figure 3C shows a large EV from filtered media that contains misfolded huntingtin cargo (green) stained with deep red membrane stain (red). Deconvolution microscopy reveals the presence of multiple GFP-72Q huntingtin^exon1^ aggregates within a single membraneous structure (Figure 3D), however, not all EVs contain huntingtin aggregates (Figure 3E). The size distribution, of the large EVs is shown in Figure 3E. These GFP-72Q huntingtin^exon1^ aggregates were found to be proteinase K sensitive as anticipated (Figure 3F).

**Figure 3.**
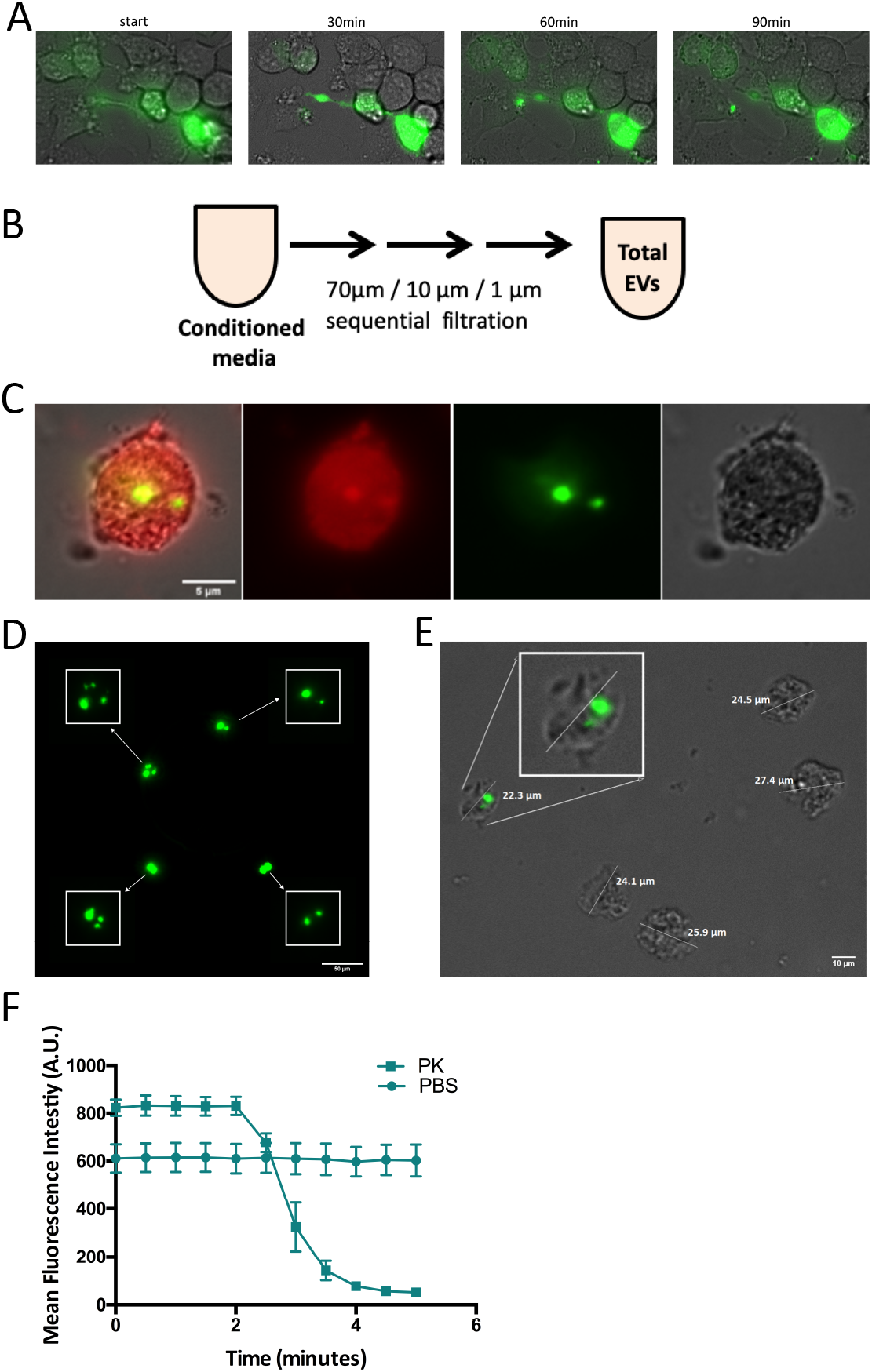
CSPα export of 15-30 μm EVs. (A) Widefield time lapse fluorescent microscopy with brightfield of EV export containing GFP-tagged huntingtin aggregates from a donor cell. (B) Experimental approach, EVs were examined following filtration. (C) Deep red membrane stain of an EV containing GFP-tagged huntingtin aggregates. (D) High-resolution, fluorescence deconvolution microscopy of GFP-tagged huntingtin aggregates located in EVs. (E) Size of EVs exported from CAD cells. (F) EVs were treated with 0.01 mg/ml proteinase K 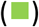 or PBS 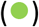 and fluorescence monitored over time.

To address whether the large vesicles contain DNA, EVs were stained with DRAQ5. Figure 4A&B shows DRAQ5 staining following the application EVs to naïve CAD cells. Cells are clearly DRAQ5 positive, while EVs containing GFP-72Q huntingtin^exon1^ do not stain with DRAQ5 indicating the EVs do not contain nuclear-like levels of DNA. Overall, CAD cells appear to be healthy following the release (Figure 3A) or application of large EVs (Figure 4). Live cell imaging following EV application to recipient cells shows that there is no dysfunction in CAD cell division in the presence of EVs containing huntingtin cargo (Figure 4E), indicating the EVs are not toxic. Direct monitoring of the 15-30μm EVs as fluorescent puncta shows that they are stable in number (Figure 4F) and fluorescence intensity (Figure 4G) and do not undergo degradation following application to recipient CAD cells. When applied to recipient cells, EVs originating from CSPα/ GFP-72Q huntingtin^exon1^ expressing cells have more aggregates (green) compared to vector/ GFP-72Q huntingtin^exon1^ control (black). CSPα-EV export of GFP-72Q huntingtin^exon1^ increases in the presence of increasing cellular CSPα expression (Figure 5A) and co-expression of GFP-72Q huntingtin^exon1^ with another control protein, a-synuclein, does not result in mutant huntingtin export. Cell viability evaluated after media collection was not influenced by CSPα expression (Figure 5B), further indicating that healthy cells export GFP-72Q huntingtin^exon1^ and export is not a consequence of cell lysis. Live cell imaging shows delivery (Figure 5C) of EVs secreted from cells expressing CSPα and labeled with deep red membrane stain. Delivery of EVs was also confirmed with ExoGlow-Protein Red and Exo Red RNA cargo labeling (data not shown). Based on these observations, it can be concluded that the huntingtin aggregates, exported in the presence of CSPα are stable and non-toxic to cells.

**Figure 4.**
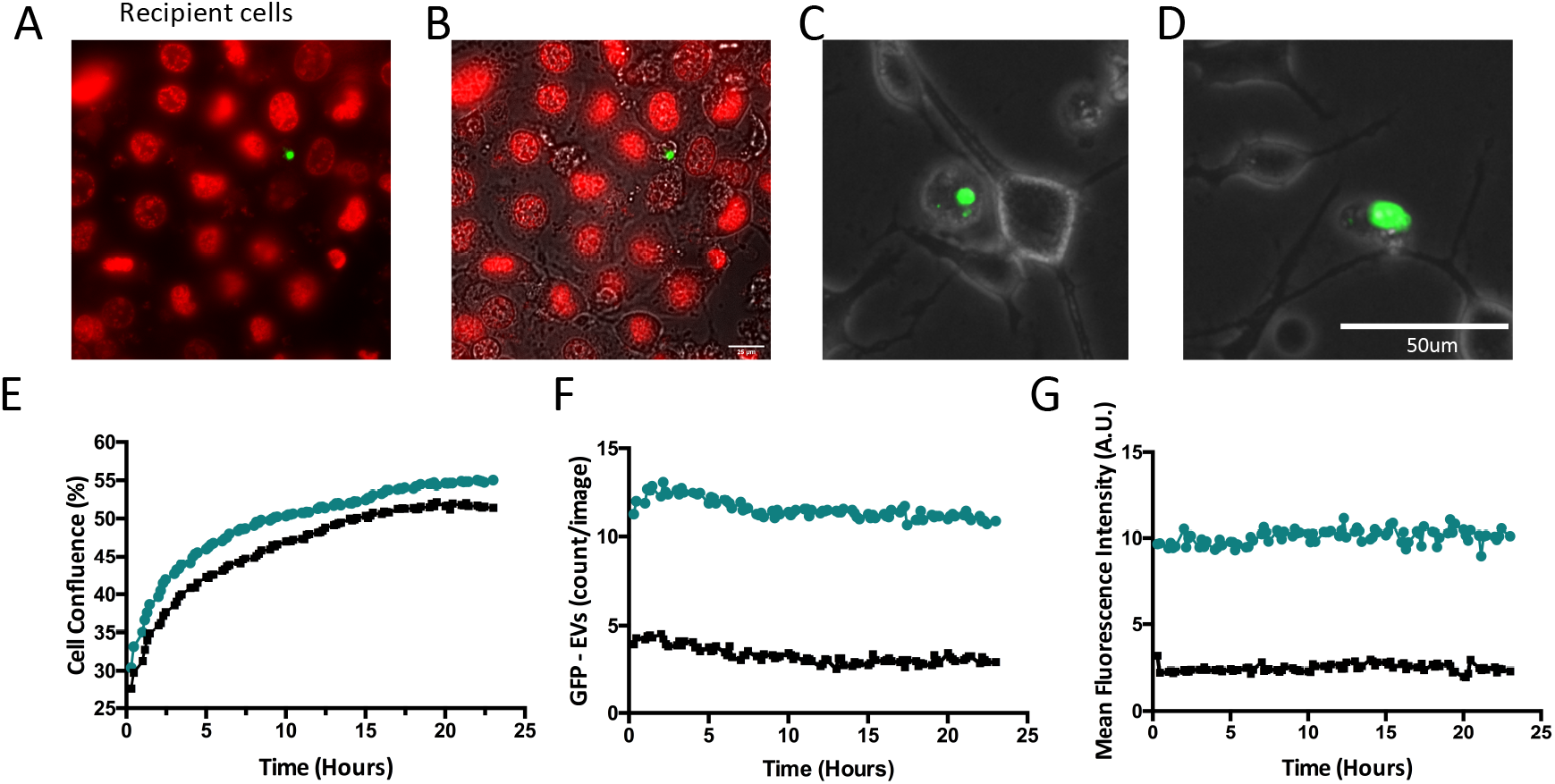
15-30 μM EVs containing GFP-tagged huntingtin do not stain with DRAQ5. EVs from CAD cells co-expressing GFP-mutant huntingtin and CSPα were applied to naïve CAD cells in 6 well plates (A) EVs containing GFP-tagged huntingtin aggregates and naïve CAD cells were stained with DRAQ5 for DNA and imaged with Widefield fluorescence microscopy. (B) With brightfield (bar = 25μm). (C&D) Representative EVOS fluorescent images of recipient cells (bar = 50μM). (E) IncuCyte live cell analysis of cell confluence following EVs application to CAD cells (Green represents EVs from cells expressing CSPα, black represents EVs from cells expressing vector (control)). (F) GFP fluorescence (puncta intensity=1, rdiameter=10 averaged over 18 images). (G) Fluorescence (relative units; mean from 18 images).

**Figure 5.**
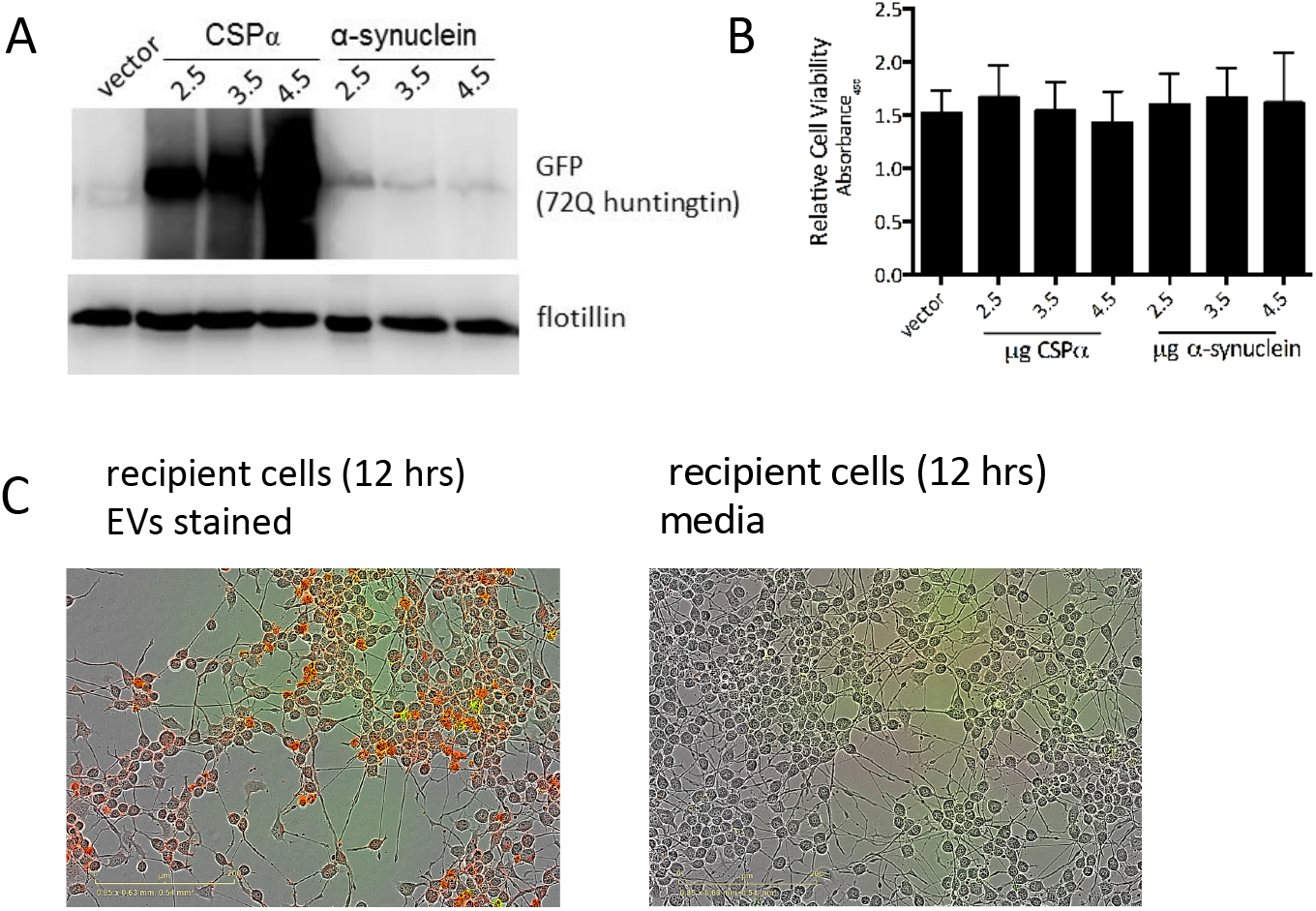
Characterization of EVs. (A) Western analysis of EVs collected from CAD cells expressing GFP-tagged 72Q Htt^exon1^ and CSPα or α-synuclein (control protein). Blots are probed for GFP and flotillin. Western blot is representative of 4 independent experiments. Flotillin is shown as a loading control. (B) Relative cell viability of CAD cells following media collection. (C) Representative incucyte images 12 hours following EV or media application to recipient cells. EVs (not enriched) were labeled with Deep Red Membrane Stain Protein Red prior to application to recipient cells. White scale bar = 50μm.

### Resveratrol Inhibits CSPα-EV Export of Mutant Huntingtin

CSPα (DnaJC5) is a type III J Protein with a well-defined domain architecture that includes an N terminal J domain and a cysteine string region (Braun et al., 1996). CSPα_L115R_ and CSPα_Δ116_ are human CSPα mutations that cause an adult onset lysosomal storage disease, ANCL (Benitez et al., 2011; Noskova et al., 2011; Velinov et al., 2012). To test the ability of CSPα_L115R_ and CSPα_Δ116_ to export mutant huntingtin in 180-240nm EVs, CSPα mutants and mutant huntingtin were transiently co-expressed in CAD cells and EVs containing GFP-tagged 72Q huntingtin^exon1^ were assessed using nanoscale flow cytometry. CSPα_L115R_ and CSPα_Δ116_ effectively mediate the export of mutant huntingtin in 180-240nm EVs, suggesting that export of aggregated proteins is not impaired in ANCL.

Resveratrol reduces export of mutant huntingtin by CSPα (Deng et al., 2017), therefore, we sought to determine whether the 180-240nm EV export was affected by resveratrol. To do so, we co-expressed GFP-tagged 72Q huntingtin^exon1^ with CSPα, CSPα_L115R_, or CSPα_Δ116_ and then treated with 50μM resveratrol 6 hours following transfection and collected EVs 24 hours following transfection for analysis. Nanoscale flow cytometry shows that resveratrol significantly reduces the CSPα, CSPα_L115R_ and CSPα_Δ116_-EV export of GFP-tagged 72Q huntingtin^exon1^ (Figure 6C). This resveratrol-decrease in export of mutant huntingtin is coupled to an increase in the overall EV export. Figure 6D shows the concentration dependency of the resveratrol reduction in secretion of the 180nm-240nm GFP labeled EV pool. The increase in EV export and decrease in GFP-72Q huntingtin^exon1^ export observed in vector-expressing cells is not significant.

**Figure 6.**
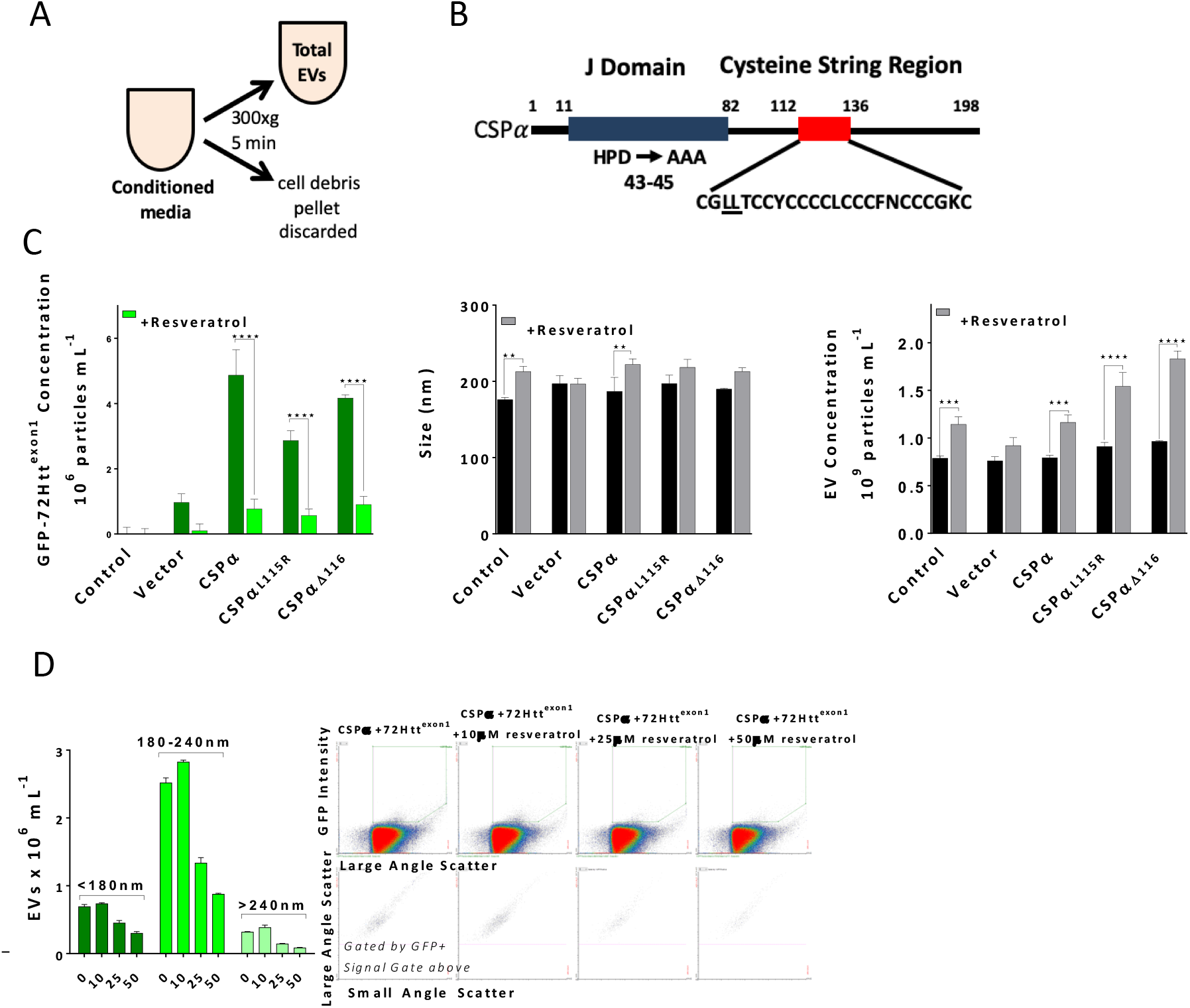
Resveratrol reduces CSPα-EV export of mutant huntingtin via 180-240nm EVs. (A) Experimental Approach, EVs were examined without enrichment. (B) Domain structure of CSPα highlighting the CSPα_HPD-AAA_, CSPα_L115R_ and CSPα_Δ116_ mutations. (C) Bar graph of the concentration of GFP-72Htt^exon1^-containing EVs collected from CAD cells expressing GFP-72Htt^exon1^ and CSPα, CSPα_L115R_, CSPα_Δ116_ or vector alone in the presence and absence of 50μM resveratrol determined by nanoscale flow cytometry. The mean size of EVs (middle panel) and the total concentration of EVs (right panel). (D) GFP-72Htt^exon1^ containing EVs flow cytometry scatter plot. (**P<0.01, ***P<0.001, ****P<0.0001)

To establish whether resveratrol influences EV cargo loading or EV export into CSPα-EVs the export of GFP huntingtin and CSPα in the presence and absence of resveratrol was evaluated by western blot. Figure 7A&B show that resveratrol reduces secretion of GFP mutant huntingtin without reducing CSPα release, suggesting that resveratrol influences loading of GFP-72Q huntingtin^exon1^ cargo into EVs without altering EV export. Cell viability following media removal is shown in Figure 7C, demonstrating that CSPα, CSPα mutants and resveratrol do not increase cell lysis and cell death. Next we evaluated the effect of CSPα mutations on EV genesis and export by examining the influence of mutant CSPα’s on EVs exported in the absence of GFP-72Q huntingtin^exon1^. The nanoparticle tracking analysis profile of the loss of function mutation, CSPα_HPD-AAA_, and the human mutations CSPα_L115R_ and CSPα_Δ116_ relative to CSPα (blue line) is shown for comparison (Figure 7D). Evaluation of total EVs within this ~30nm-1000nm range shows lower amounts of the minor peaks at 200nm and 370nm by CSPα_L115R_ and CSPα_Δ116_ compared to CSPα. No difference in mean size of EVs was observed as a result of CSPα mutations (Figure 7E) and the distribution of <180nm, 180-240nm, >240nm of EVs was the same in the presence of CSPα mutants (data not shown). Taken together, our data indicate that resveratrol reduces export of mutant huntingtin without blocking EV export.

**Figure 7.**
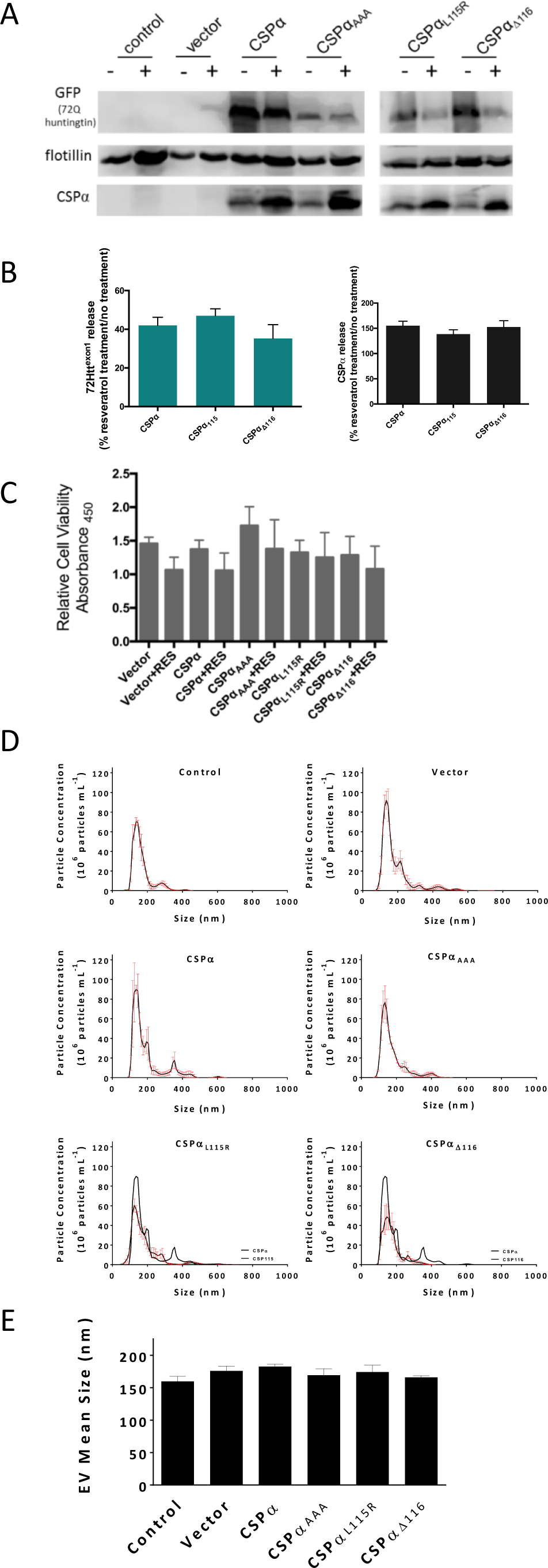
Influence of CSPα mutants on EV export. (A) Western analysis of EVs collected from CAD cells expressing GFP-72Htt^exon1^ and CSPα, CSPα_AAA_, CSPα_L115R_, CSPα_Δ116_ or vector in the presence (+) and absence (-) of 50 μM resveratrol. Western blots are probed for GFP, flotillin and CSPα. (B) Western blot quantification (C) Relative cell viability of CAD cells post media collection. (D) NTA analysis of total EVs exported from CAD cells expressing CSP? mutants (but not GFP-72Htt^exon1^); Mean histogram data for CSPα is overlaid in blue for comparison. (E) Mean size of total EVs exported from CAD cells.

We then asked whether resveratrol also reduces mutant huntingtin export in 15-30μm EVs. EVs were collected from CAD cells 48 hours following expression of GFP-72Qhuntingtin^exon1^ with CSPα and processed by sequential filtration and applied to recipient cells. Fewer large EVs with aggregates originated from CSPα/ GFP-72Qhuntingtin^exon1^expressing cells in the presence (black) compared to the absence of resveratrol (green). Neither the number of EVs carrying GFP-72Qhuntingtin^exon1^ cargo or the GFP fluorescence diminishes over time, demonstrating the stability of exported mutant huntingtin (Figure 8B). Figure 8C shows recipient cells after application of EVs from CSPα/GFP-72Qhuntingtin^exon1^ expressing cells treated with or without resveratrol. Although lower in abundance, the 15-30μm EVs are clearly detectable following resveratrol treatment. Figure 8D compares images of large EVs containing misfolded huntingtin from filtered media collected from CSPα-expressing cells in the absence and presence of resveratrol.

**Figure 8.**
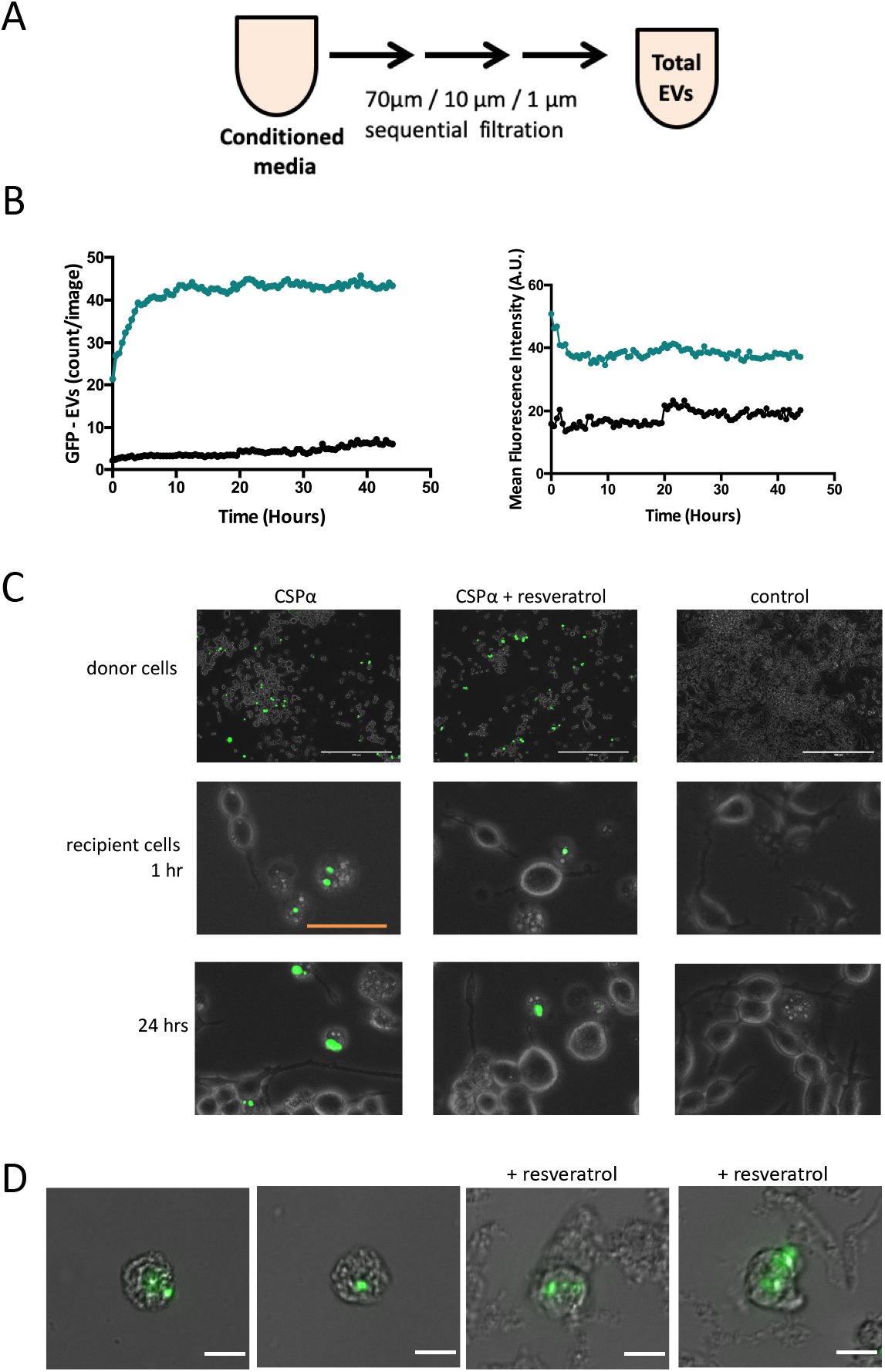
Resveratrol reduces CSPα-EV Export of mutant huntingtin by 15-30nm EVs. (A) Experimental Approach, EVs were examined following filtration. (B) EVs from CAD cells coexpressing GFP-mutant huntingtin and CSPα in the absence (green) and presence of resveratrol (black) were applied to naïve CAD cells in 12 well plates. IncuCyte live cell analysis of GFP EVs (left panel) puncta intensity=1, rdiameter=10 and mean fluorescence averaged over 9 images and (right panel) fluorescence (relative units; mean from 9 images). (C) GFP-72Htt^exon1^ aggregates in donor CAD cells at the time of media collection and GFP-72Htt^exon1^ aggregates in recipient cells at 1 and 24 hrs. (D) EVs containing GFP-tagged huntingtin aggregates and naïve CAD cells were imaged with Widefield fluorescence microscopy.

The export of EVs carrying mutant huntingtin cargo by CSPα is dependent on the histidine, proline, aspartate (HPD) motif, located within the J domain of CSPα (Deng et al., 2017). CSPα_HPD-AAA_ is a loss-of-function mutant that does not activate Hsp70ATPase and we surmised would not facilitate export of GFP-72Qhuntingtin^exon1^ in 15-30μm EVs. Figure 9A shows a manual count of large mutant huntingtin-containing EVs obtained from cells expressing vector (control), CSPα and CSPα_HPD-AAA_ 24 hours following application to recipient cells that is consistent with the live cell imaging count. GFP-72Qhuntingtin^exon1^ in EVs was unaffected by direct application of resveratrol to already exported EVs, indicating that resveratrol has a preEV export mode of action (Figure 9B). Finally, we asked whether CSPα_L115R_ and CSPα_Δ116_ stimulate release of GFP-72Qhuntingtin^exon1^ in the large 15-30μm EVs. CSPα_L115R_ and CSPα_Δ116_ support robust export of GFP-72Qhuntingtin^exon1^ in 15-30μm EVs (Figure 9C&D) and export is blocked in the presence of 50μm resveratrol, indicating that residues 115 and 116 are not essential for export.

**Figure 9.**
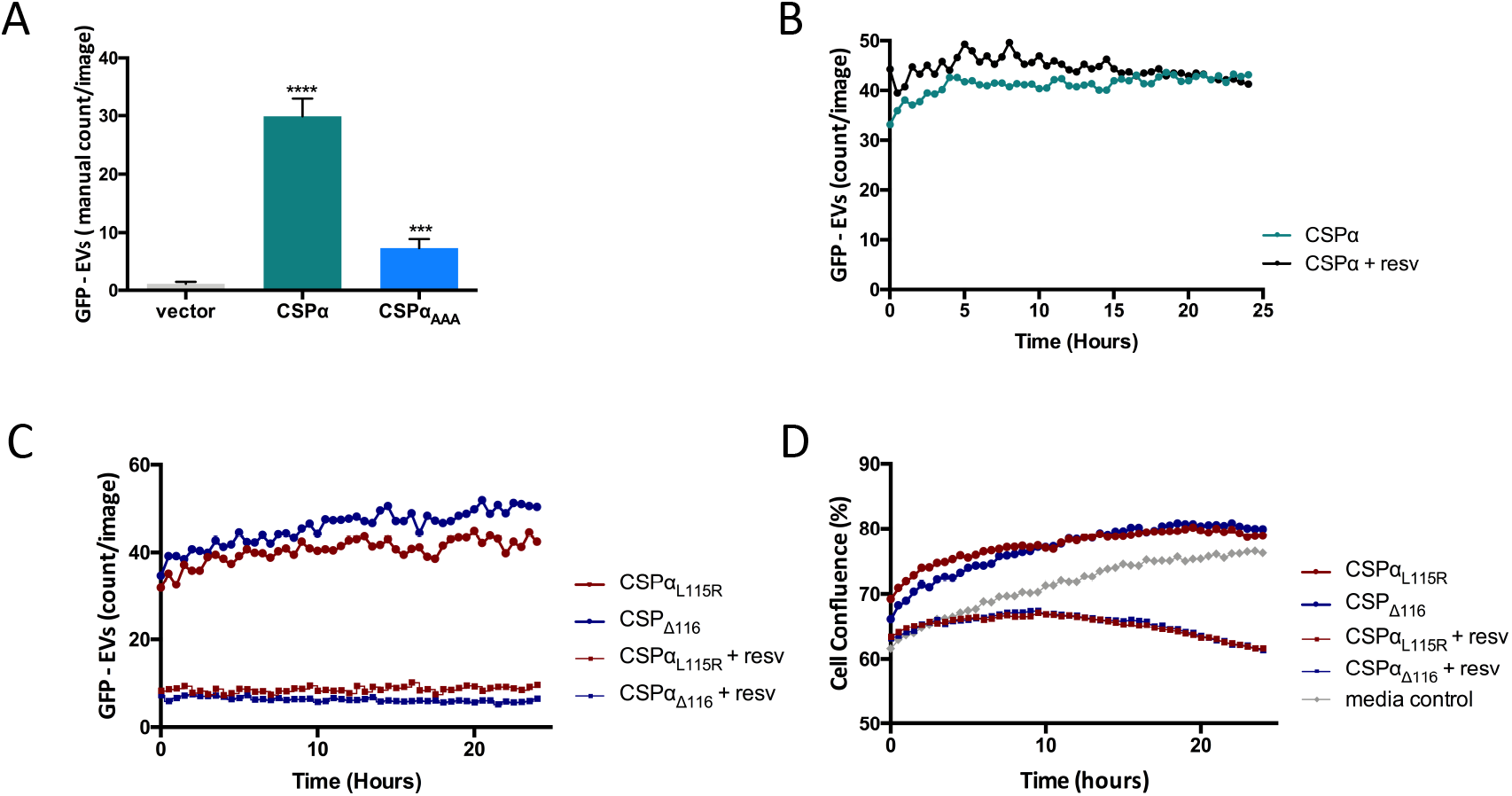
Resveratrol reduces EV Export of mutant huntingtin by CSPα and CSPα mutants. (A) Manual count of GFP-EVs. EVs from cells expressing GFP-mutant huntingtin and the indicated proteins were applied to recipient cells at 24 hours (n=10, ***P<0.001, ****P<0.0001). (B-D) IncuCyte live cell analysis of GFP EVs following application to naïve cells in 12 well plates, puncta intensity=1, rdiameter=10 and mean fluorescence averaged over 9 images. (B) Direct application of 50μm resveratrol with EVs from cells expressing GFP-mutant huntingtin and CSPα (donor cells not treated with resveratrol). (C) EVs from cells expressing GFP-mutant huntingtin and CSPα mutants, +/- 50μm resveratrol. (D) IncuCyte live cell analysis of cell confluence following application of EVs.

Collectively, our results show that CSPα-mediates export of mutant huntingtin through two subpopulations of EVs, sized at 180-240nm and 15-30μm and that resveratrol blocks export of mutant huntingtin through both EV routes. The large EVs contain multiple mutant huntingtin aggregates which are pliable and sensitive to proteinase K degradation, yet stable when applied to recipient cells. GFP-tagged 72Qhuntingtin^exon1^ export in EVs is dependent on Histidine, Proline Aspartate (HPD) motif, located within the J domain of CSPα, however disease related mutations do not impact export capacity.

## Discussion

Multiple pathways for the cell-to-cell transfer of misfolded proteins have emerged in addition to 180-240nm and 15-30μm EVs, including tunneling nanotubes, (4μm) exofers and free floating proteins (Costanzo et al., 2013; Melentijevic et al., 2017). These systems operate to eliminate misfolded proteins in conjunction with the universal systems, proteosomal degradation, autophagy-mediated protein turnover and chaperone-assisted protein folding. Our data demonstrate that the co-chaperone CSPα, facilitates mutant huntingtin export in 180-240nm and 15-30μm EVs. Although, cells release EVs that are markedly heterogenous in size, mutant huntingtin is found in only two sizes of EVs: 180-240nm and 15-30μm. We speculate that export of mutant huntingtin is a mechanism to maintain proteostasis in donor cells and dispose of harmful misfolded proteins to cells with greater clearance capacity for toxic protein aggregates. This would be particularly relevant in neuronal synapses where the basic proteostatic machinery is limited. Identification of two separate EV export pathways argues for the importance of export in managing misfolded protein levels, however the role these two export pathways play in neurodegenerative disease progression remains under investigation.

Extensive evidence supports a role of CSPα, in neural quality control mechanisms. CSPα knock-out mice exhibit fulminant neurodegeneration and have a reduced lifespan with no mice surviving beyond 3 months (Fernandez-Chacon et al., 2004). Loss-of-function CSPα *Drosophila* mutants demonstrate uncoordinated movements, temperature-sensitive paralysis and early lethality (Zinsmaier et al., 1994). And, in *C elegans*, CSPα null mutants display neurodegeneration and reduced lifespan (Kashyap et al., 2014). Compelling evidence has been presented that several disease-causing proteins are exported through the CSPα export pathway We have shown the CSPα EV export pathway exports SOD-1^G93A^ and 72Q huntingtin^exon1^ (Deng et al., 2017). Fontaine and colleagues have demonstrated that CSPα increases secretion of TDP-43, α-synuclein and tau from HEK293 cells (Fontaine et al., 2016). Furthermore, Ye and colleagues have shown that CSPα facilitates export of, TDP-43, a-synuclein and tau from HEK and COS7 cells(Lee et al., 2018; Xu et al., 2018). Not surprisingly, CSPα dysfunction has been implicated in several neurodegenerative disorders in addition to ANCL, including Alzheimer’s, Parkinson’s and Huntington’s disease (Chandra et al., 2005; Donnelier et al., 2015; Henderson et al., 2016; Miller et al., 2003; Tiwari et al., 2015; Zhang et al., 2012). A *c. elegans* screen identified the polyphenol, resveratrol, to ameliorate the reduced life span of CSPα mutants suggesting a role in proteostasis (Kashyap et al., 2014a). Resveratrol is an established multitarget directed compound, and how the longer life span relates to a resveratrol-mediated reduction in EV export of mutant huntingtin remains to be determined.

Neural export of misfolded proteins offers many advantages such as maintenance of parent cell protein quality control but also many dangers if unregulated. Our results illustrate a double EV export system that removes toxic huntingtin from cells and link the molecular cochaperone CSPα to EV genesis and export. We speculate that the cell-to-cell transfer of toxic huntingtin becomes progressively dysregulated during Huntington’s disease progression. The interplay between EV export and cellular degradative systems remain under investigation.

## Supplementary Figures

**Figure S1.**
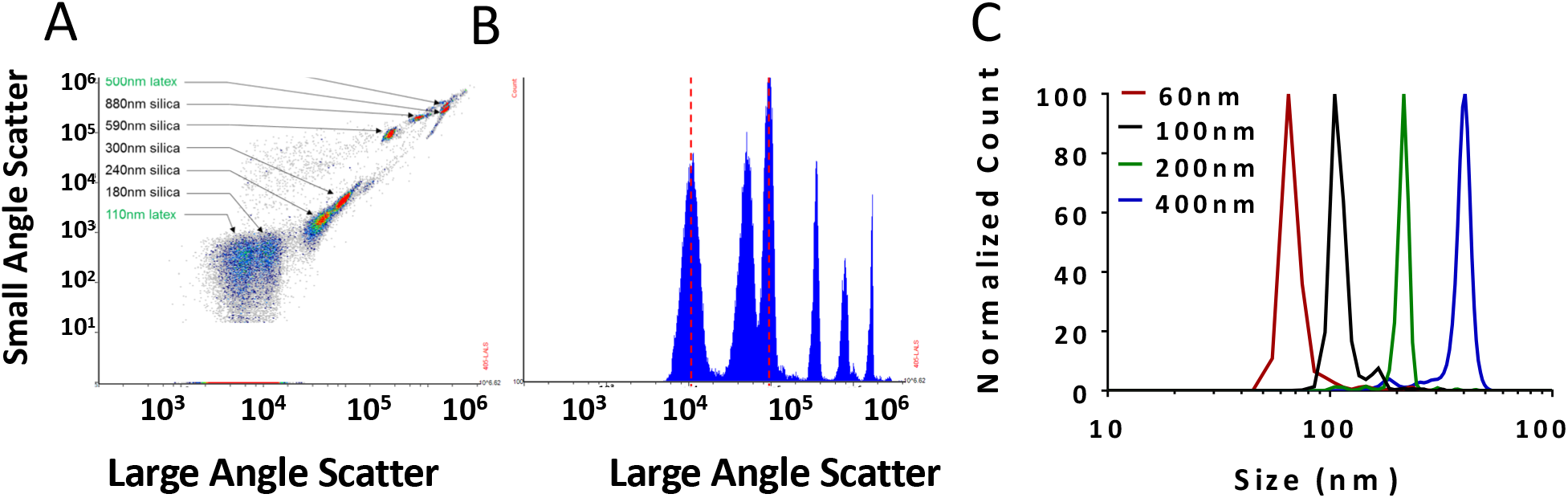
(A) Scatter of nanoscale flow cytometry size standards including both fluorescent latex and non-fluorescent silica standards ranging in size from 110nm – 1300nm. (B) Red lines illustrate the division of size ranges of detected EVs described in the text to carry GFP-tagged 72Q Htt^exon1^ cargo. (C) NTA size standards (mean) included 60nm, 200nm and 400nm polystyrene NIST size standards as well as a 100nm silica size standards.

**Figure S2.**
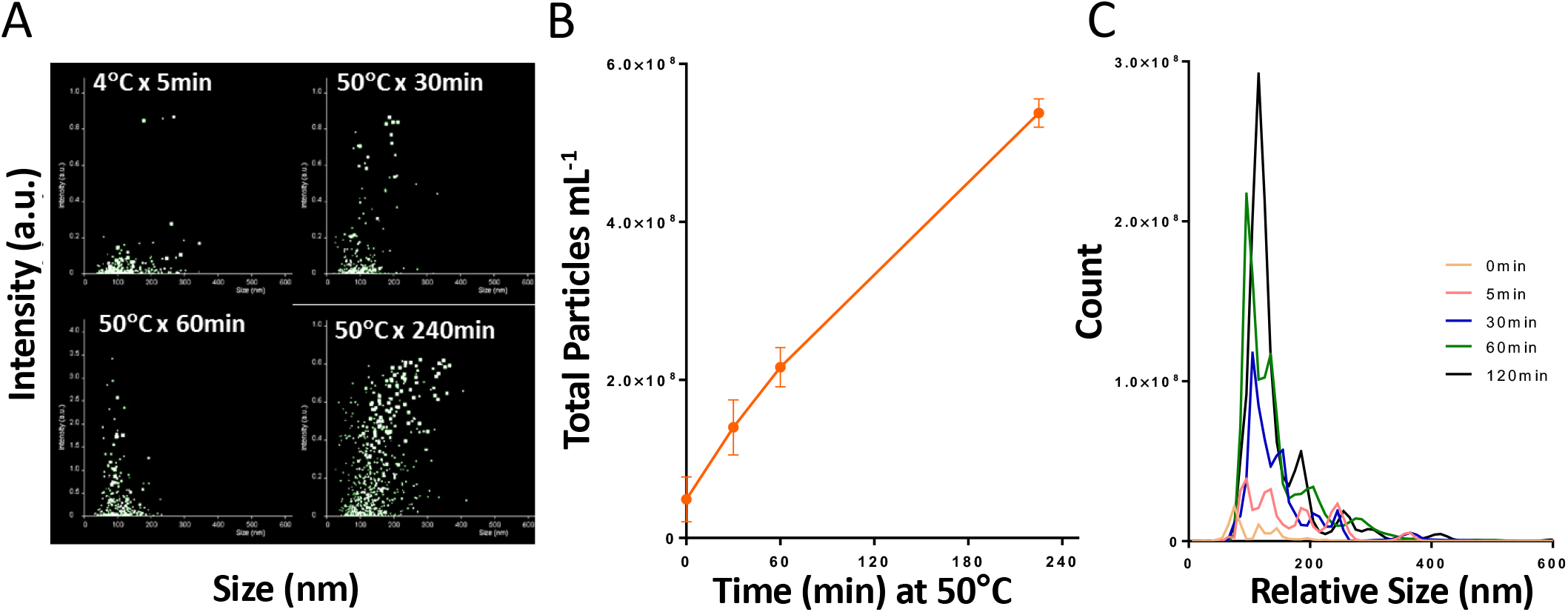
(A) NTA analysis of anti-CD36FITC antibody aggregation. (B) Line graph of the time dependent aggregation of anti-CD36-PE antibody (C) NTA profiles of anti-CD36-FITC (or PE) antibody aggregation.

## Materials and Methods

### EV collection from CAD cells

For EV production, CAD cells (catecholaminergic derived CNS cells) were seeded in 10 cm culture dishes in Dulbecco’s Modified Eagle’s Medium (DMEM-F12 Gibco Thermo Fisher Scientific), supplemented with 10% Foetal Bovine Serum (FBS, Gibco Thermo Fisher Scientific), and 1% penicillin (100U/ml) and streptomycin (100ug/ml) (P/S, Gibco Thermo Fisher Scientific) and maintained at 37C, 5% CO2 atmosphere. After 24 hours cells were transfected with indicated plasmids using Lipofectamine 3000 (Invitrogen) in Opti-MEM™ medium. 6 hours after transfection, medium was changed to serum free Dulbecco’s Modified Eagle’s Medium (DMEM Gibco Thermo Fisher Scientific), supplemented with 1% penicillin (100U/ml) and streptomycin (100ug/ml) (P/S, Thermo Fisher Scientific). Conditioned medium was collected 48 hours after transfection and spun at 300Xg for 5 min to remove cell debris and larger particles and then evaluated by nanoscale flow cytometry analysis and live cell imaging. Following media collection, cell viability was determined utilizing an XTT assay (New England Biolabs). Similar results were obtained utilizing calcium phosphate transfection methods. Where indicated media was processed by sequential filtration through 70μm (nylon) filters followed by 10μm/1μm (PET; Polyethylenterephthalat) filters (pluriSelect) with negative pressure applied via a syringe (Figures 3, 4 and 8).

### Immunoblotting

For western analysis following the 300Xg spin EVs were subjected to exoquick precipitation solution (SBI) and solubilized in sample buffer. Proteins were separated by SDS-PAGE and electrotransferred from polyacrylamide gels to nitrocellulose membrane (0.2μm pore size). Membranes were blocked in tris-buffered saline (TBS) containing 0.1 % Tween 20, 1 % BSA and then incubated with primary antibody overnight at 4°C. The membranes were washed and incubated with horseradish peroxidase-coupled secondary antibody for ~2 hours at room temperature. Bound antibodies on the membranes were detected by incubation with Pierce chemiluminescent reagent and exposure to Cdigit, LiCor (Mandel). The chemiluminescent signals were quantified using image studio digits software (Mandel).

### Plasmids

cDNAs encoding for CSPα, CSPα mutants and α-synuclein were expressed in the plasmid myc-pCMV. GFP-72Q huntingtin^exon1^ was expressed in the plasmid pcDNA3.1. All amplified regions of all plasmids were sequenced to ensure the absence of any undesired mutations.

### Fluorescence Imaging

Nikon widefield images were acquired at room temperature on a Nikon Ti Eclipse widefield microscope equipped with a Hamamatsu Orca flash 4.0 v2 sCMOS 16-bit camera using NIS-Elements AR v5.00.00 64-bit software. Images were captured using either a 40x Plan Apo λ/0.95 numerical aperture (NA) objective or a 60x Plan Apo λ/1.4 NA oil objective.

Incucyte images were acquired on an Essen BioScience IncuCyte Zoom microscope using IncuCyte Zoom v2018A software. The IncuCyte Zoom microscope was in a humidified incubator at 37°C with 5% CO_2_. Nine images per well of either six-well plates (Figure 4) or twelve-well plates (Figures 8 and 9) were captured every 15 min for 24 hour using a 20x Plan Fluor/0.4 NA objective.

Analysis was performed using IncuCyte Zoom v2018A software. Analysis masks were created to determine confluency (confluence mask) and presence of green objects (green object mask). These masks were applied to all images in a data set.

EVOS FL auto images of live donor cells were taken at the time of media collection. Images of recipient cells were taken between 1-24 hours. The intensity of the GFP-72Q Htt^exon1^ varied among cells and we used the single slider until the on-screen brightness of the lowest intensity aggregate was satisfactory and used this setting to capture images in high-quality mode. For images of Cell Mask Deep Red Plasma Membrane Stain Red-labeled EVs, EVs were labeled as per manufacturers instructions (Systems Biosciences), washed two times in media and applied to recipient CAD cells for 12hrs.

### Nanoparticle Tracking Analysis

Particle size and concentration of the samples were determined via nanoparticle tracking analysis (NTA) using Nanosight LM10 equipped with 405nm laser, 60mW, software version 3.00064). Briefly, samples were diluted 25 fold using 0.2x phosphate buffered saline that had been filtered twice through 0.2micron filter and analyzed in triplicate for 60 seconds per replicate with a count range of 20-100. All samples were analyzed particles per frame. A variety of NIST (Thermo Scientific 3000 Series) standards were analyzed each day prior to sample analysis. The system was cleaned between each sample and checked for any sample carryover using the PBS diluent.

### Nanoscale Flow Cytometry

Particle size and concentration of the samples were determined via nanoscale flow cytometry using Apogee A50 flow cytometry platform. Samples were similarly prepared for NTA and Nanoscale Flow Cytometry. Light scatter was provided using the 405nm laser (75mW); GFP signal was generated using the 488nm laser (50mW, 535/35) and far red signal was generated using the 630nm laser (75mW, 680/35). All samples were analyzed for 60 seconds. Optimization was performed to insure single EVs were being analyzed and single events were triggered by light scatter only. The system was cleaned each day prior to sample analysis and a variety of silica and polystyrene standards (Apogee 1493 standards) processed for instrument set up and QC. The silica standards were used to assess the relative size range of EVs (Supplementary Figure 1).

To demonstrate that the GFP-72Q Htt^exon1^ signal was associated with bona fide vesicles and not simply protein aggregates, samples were first mixed with Cell Mask Deep Red plasma membrane stain (Thermo Fisher Scientific; C10046), final concentration of 0.1X), incubated for 30 minutes at 37°C and then diluted in PBS. For comparison, palmitoylated-GFP positive EVs were obtained from the conditioned media of a PC3 prostrate cancer cell line; the membrane of these EVs contain the FP proteins. 10μl of anti-CD-FITC antibody was incubated at 50°C for increasing periods of time to generate non-membrane, protein aggregates. The protein aggregate controls as well as the PC PALM-GFP EV controls were similarly stained with Deep Red cell mask. Aggregate concentration was analyzed by NTA and uptake of the membrane dye was measured using nanoscale flow cytometry. For these experiments, single particle detection was triggered using positive GFP/FITC fluorescence and the associated far red signal analyzed.

### Statistical Analysis

All data were graphed and statistically analyzed using GraphPad Prism version 6.01 for Windows, GraphPad Software, La Jolla California USA, www.graphpad.com. Statistics included One-Way and Two Way ANOVA with either Tukey’s or Dunnett’s post test analysis if initial ANOVA was statistically significant (p<0.05; stars indicate significance ★ p<0.05, ★★ p<0.01, ★★★ p<0.001, ★★★★ p<0.0001). All values are presented as the mean ±SEM where appropriate, otherwise the SD is presented as indicated.

## Acknowledgements

The work was supported by a grant from the Alzheimer Society of Alberta and Northwest Territories, the Alberta Prion Research Institute and National Research Council of Canada. The authors would like to express their gratitude to Dr. Frank Visser for technical support. Perrin Beatty assisted with figure preparation. We acknowledge the work done using the Live Cell Imaging Facility at the University of Calgary.

## Author Contributions

J.E.A.B. conceived the project. D.P. and J.L. designed and interpreted all NTA and NFC experiments. J.E.A.B designed and interpreted all WB and FI experiments. J.D. provided technical assistance. J.E.A.B. D.P. and J.L. wrote the manuscript.

